# Key roles of microbial sulfur and ammonium oxidizers for the coastal seafloor ecological state

**DOI:** 10.1101/2024.09.19.613430

**Authors:** Tonje Nilsen, Ragnhild Pettersen, Nigel Brian Keeley, Jessica Louise Ray, Sanna Majaneva, Morten Stokkan, Anja Hervik, Inga Leena Angell, Lars Gustav Snipen, Maud Ødegaard Sundt, Knut Rudi

## Abstract

Recent evidence suggests that there is a major switch in coastal seafloor microbial ecology already at a mildly deteriorated macrofaunal state. This knowledge is of critical value in the management and conservation of the coastal seafloor. We therefore aimed to determine the relationships between seafloor microbiota and macrofauna on a regional scale. We compared prokaryote, macrofauna, chemical, and geographical data from 1,546 seafloor samples which varied in their exposure to aquaculture activities along the Norwegian and Icelandic coasts. We found that the seafloor samples contained either a sulfur oxidizer network (42.4% of samples, n=656), or an ammonium oxidizer network of microbes (44.0% of samples, n=681). Very few samples contained neither network (9.8% of samples, n=151), or both (3.8% of samples, n=58). Samples with a sulfur oxidizer network had a tenfold higher risk of macrofauna loss (odds ratios, 95% CI: 9.5 to 15.6), while those with an ammonium oxidizer network had a tenfold lower risk (95% CI: 0.068 to 0.11). The sulfur oxidizer network was negatively correlated to distance from Norwegian aquaculture sites (Spearman rho = −0.42, p < 0.01), and was present in all Icelandic samples (n=274). The ammonium oxidizer network was absent from Icelandic samples, and positively correlated to distance from Norwegian aquaculture sites (Spearman rho = 0.67, p < 0.01). Based on 357 high-quality metagenome-assembled genomes (MAGs), we found that the main metabolic process for the ammonium oxidizer network was cobalamin-dependent, while the sulfur oxidizer network was associated with both ammonium retention and sulfur metabolism. In conclusion, our findings highlight the critical roles of sulfur and ammonium oxidizers in mild macrofauna deterioration, which should be included as an essential part of seafloor surveillance.

## INTRODUCTION

The coastal seafloor represents one of the richest and most productive marine ecosystems ^1^. The seafloor is an integral component of ocean ecosystems, providing a habitat for biological diversity as well as essential services such as nutrient recycling and production of trace elements ^2^. Despite the large geographical area encompassed by the coastal seafloor, the factors influencing seafloor biodiversity and ecosystem function are poorly understood ^3,4^. This lack of knowledge is particularly critical, as the coastal seafloor can be a recipient for anthropogenic waste, such as that from aquaculture farms ^5^, sewage ^6^ and industry ^7^.

In a geographically restricted study, we have previously shown that the seafloor microbiota in the northern part of Norway demonstrated bimodal diversity distribution, with low-diversity prokaryote communities associated with the sulfur-oxidizing genus *Sulfurovum* ^8^. In the same study, we showed that high-diversity prokaryote communities were associated with the ammonium oxidizing archaeon *Nitrosopumilus* ^8^. This was a surprising discovery as the main diversity transition from high to low diversity occurred in an area of oxygenated seafloor. *Nitrosopumilus* is one of the most abundant microorganisms in the ocean ^9,10^, while its role in supporting biodiversity is unknown. *Sulfurovum* has primarily been associated with geothermal-, rather than human, activities ^11^, although there are some reports of noteworthy abundances related to dredged sediments and fish farms ^12,13^. It is therefore unclear whether the bimodal *Nitrosopumilus*-*Sulfurovum* distributions observed for Northern Norwegian sediments ^8^ are representative at a regional scale, or how the microbial composition and functional potential relates to macrofauna ecological status.

Most knowledge about the effect of human activity on seafloor diversity is derived from impact studies of aquaculture ^5,14^. To facilitate comparisons across geographic regions, various diversity indices have been developed. Some indices utilize eDNA-derived microorganism counts ^15^ in a manner comparable to the well-established macrofauna indices ^16^, wherein organisms are categorized into Eco-Groups based on their relative sensitivity to waste. The macrofaunal indices, however, are generally not harmonized between countries ^17^. In Norway and Iceland, the most widely used macrofaunal index is the normalized Ecological Quality Ratio (nEQR) ^18^. nEQR is a measure of the level of ecological status that integrates information about diversity and key macrofauna taxa. The nEQR index ranges from 0 to 1, with 0 to 0.2 being severely deteriorated, 0.2 to 0.4 deteriorated, 0.4 to 0.6 moderately deteriorated, 0.6 to 0.8 mildly deteriorated, and 0.8 to 1 representing the macrofaunal natural state ^19^.

For highly deteriorated macrofauna, it is well established that anoxia and sulfides are drivers of deterioration ^20^. We lack, however, a good understanding of mechanisms underlying the low- to non-deteriorated macrofauna. Within benthic assemblages, it is likely that microorganisms provide essential nutrients or other exogenous compounds required by macrofauna ^21,22^. Alterations of key microbial service providers could therefore lead to major unforeseen effects on the macrofauna ^23^.

This work aimed to identify prokaryotic taxa and processes associated with the macrofaunal ecological status at a regional scale, focusing on samples with high ecological status. This was achieved through large scale comparative studies, including prokaryote metabarcoding, macrofaunal assessment, metagenomic analyses, and a range of chemical measurements of more than 1,500 samples along the Norwegian and Icelandic coasts, with varying exposure to impact from aquaculture activities.

## RESULTS

A total of 1,546 sediment samples from Norwegian (41 locations) and Icelandic (9 locations) coastal regions were included in this study, with the majority covering aquaculture farm facilities (Suppl. Metadata; Suppl Table 1). We identified a total of 49,543 prokaryote OTUs (clustering at 95 % similarity) representing 2,387 genera and 61 phyla (Suppl. Taxonomy).

### Correlation networks reveal strong negative correlation between Nitrosopumilus and Sulfurovum

To determine potential microbial interactions, we investigated the pairwise correlation structure for all the prokaryote genera identified. In total, we identified 174,743 moderate correlations with Spearman rho > 0.3, and 28,557 with rho < −0.3. More than 200 genera showed over 500 positive correlations, while only one genus, *Sulfurovum*, showed more than 500 negative correlations. The overall strongest pairwise negative correlation was identified between the genera *Nitrosopumilus* and *Sulfurovum* (Spearman rho=-0.79, p<0.0005). This negative correlation was therefore used as a stem for the other genera in defining correlation patterns.

In total, 25.2 % of the genera showed separate correlation patterns for *Nitrosopumilus* and *Sulfurovum.* A large sub-network of 516 genera showed both moderate positive correlation with *Nitrosopumilus* (Spearman rho > 0.3) and moderate negative correlation with *Sulfurovum* (Spearman rho < −0.3), while 79 genera showed moderate positive correlation with *Sulfurovum* and moderate negative correlations with *Nitrosopumilus*. We therefore defined the *Nitrosopumilus* and *Sulfurovum* networks, representing the genera showing positive and negative correlation to *Nitrosopumilus* and *Sulfurovum*, respectively (Fig. 1).

**Figure 1.**
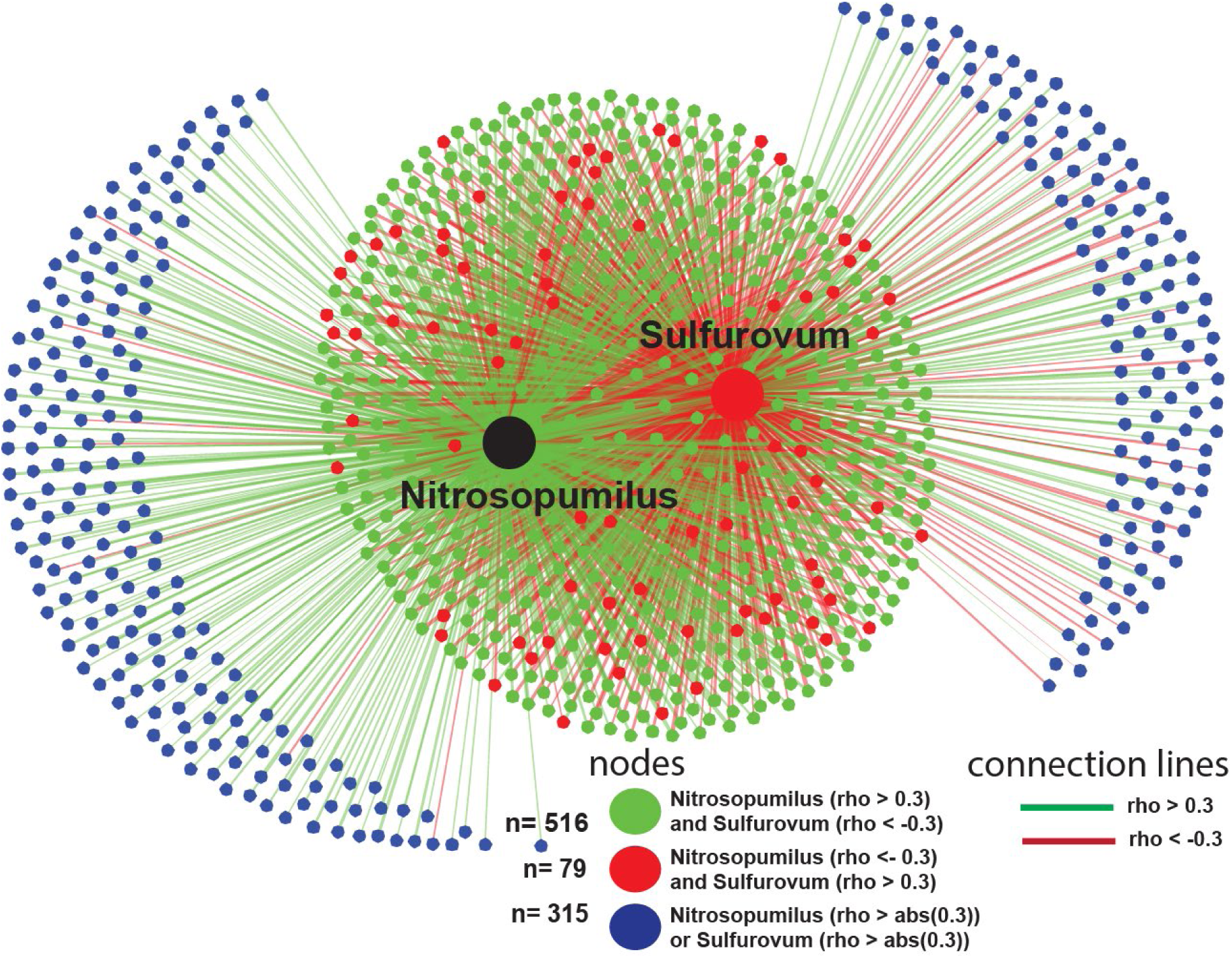
Correlation networks between *Sulfurovum* and *Nitrosopumilus*. The correlation network for *Nitrosopumilus* and *Sulfurovum* is illustrated by genera showing positive correlations with *Nitrosopumilus* and negative correlations with *Sulfurovum.* Each node represents a separate genus, while the genus *Nitrosopumilus* is highlighted in black, and the genus *Sulfurovum* in red. The color-code for the nodess represents the correlation patterns, as described in the figure. The color code for the connecting lines represents the direction correlation between the given pairs, with green lines representing Spearman rho > 0.3, while red lines represent rho < −0.3.

### The macrofaunal ecological status can be predicted based on the *Nitrosopumilus –* and *Sulfurovum* networks

Using cross-validated generalized additive models (GAM) based on the *Nitrosopumilus* and *Sulfurovum* network relative abundances, the predicted nEQR macrofauna ecological state correlated very well with the observed values (Fig. 2A). In the model, the relative abundance of both the *Sulfurovum* and *Nitrosopumilus* networks showed non-linear partial dependence to the nEQR values (Fig. 2 B and C). For the *Sulfurovum* network, there was a negative linear trend in the partial dependence to a nEQR value of about 0.4 (Fig. 2B). The relationship between nEQR and the relative content of the *Nitrosopumilus* network shows a threshold at about 0.2, as visualized in the partial dependence plot (Fig. 2C). Depth and distance from aquaculture sites showed low influence in the model, as indicated by the flat curves for the partial dependencies (Fig. 2D and E).

**Figure 2.**
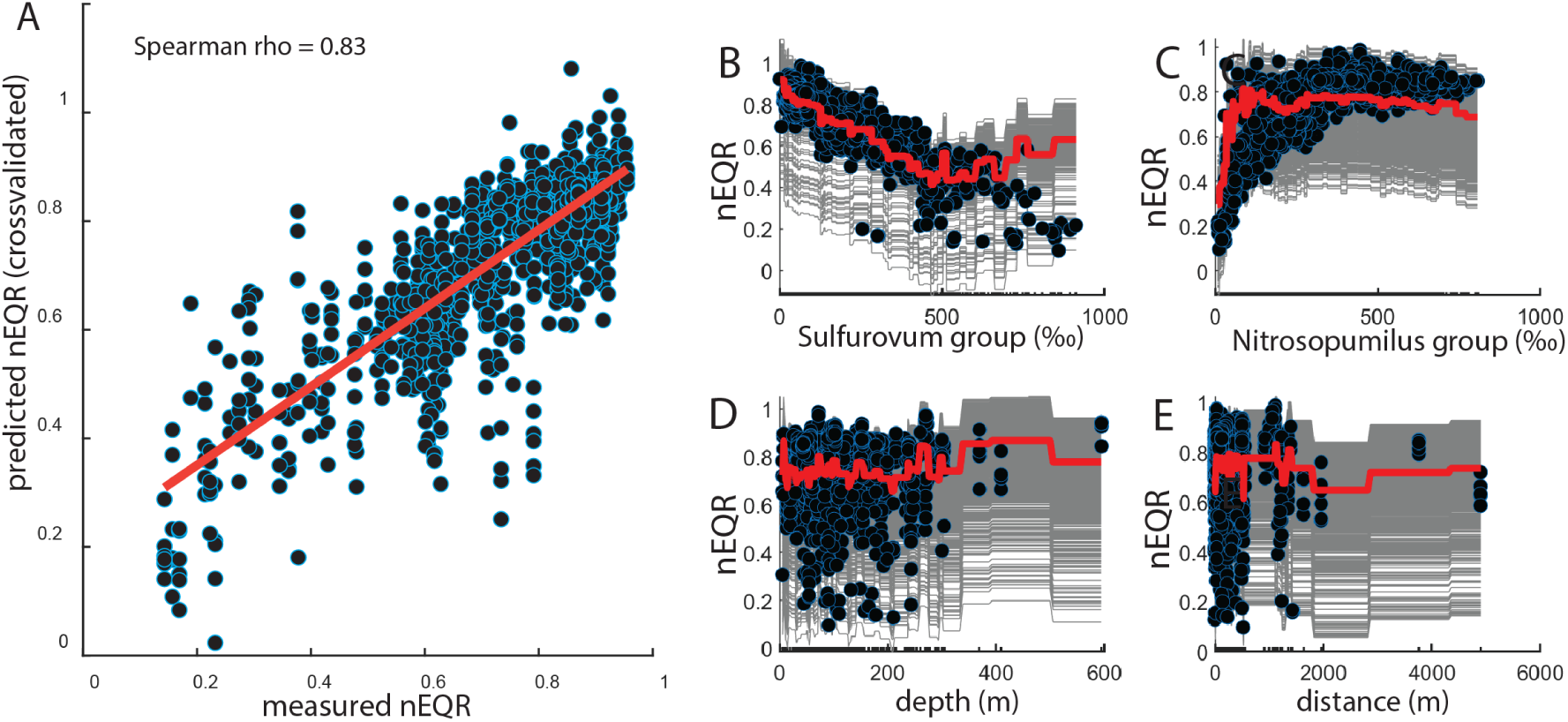
Model for the association between macrofauna and microbiota. (A) Correlation between measured nEQR, and the cross-validated prediction using the GAM model. (B to D) The dots represent the actual measurements for each of the four components in the GAM model. Individual Conditional Expectation plots are presented as grey lines for each of the measurements, while the thick red line represent the partial dependence in the GAM model.

The underlying direct associations are shown in Suppl. Fig. 1.

### The *Nitrosopumilus* and *Sulfurovum* networks showed opposite associations to chemical and physiochemical composition

The chemical measurements unveiled several strong correlations with prokaryote community composition (Fig. 3). The *Sulfurovum* network was weakly positively correlated, while the *Nitrosopumilus* network was weakly negatively correlated, to oxygen saturation above 80% (Fig. 3A). The *Nitrosopumilus* network showed a strong negative correlation with pH up to 7.9, with a switch to a strong positive correlation to pH greater than 7.9 (Fig 3B), while the *Sulfurovum* network showed a positive correlation to pH across the observed pH scale (Fig. 3B). For redox potential, there was no clear correlation of either network (Fig. 3C). There was a slight negative correlation between the *Nitrosopumilus* network and total nitrogen for values < 3.5 mg/kg, above which a positive correlation was observed. The *Sulfurovum* network demonstrated the opposite correlative trend with total nitrogen (Fig. 3D). A strong negative correlation to organic carbon was shown for the *Nitrosopumilus* network, while the *Sulfurovum* network showed a positive correlation (Fig. 3E). At a ratio between organic carbon and nitrogen below 10, the *Nitrosopumilus* network was overrepresented in comparison to the *Sulfurovum* network (Fig. 3F). The *Nitrosopumilus* network had a strong positive correlation to pelite, while the *Sulfurovum* network was negatively correlated (Fig. 3G). Furthermore, there was a positive correlation between zinc and the *Nitrosopumilus* network, while correlation to zinc was negative for the *Sulfurovum* network (Fig. 3H). Finally, copper showed a multimodal distribution, with a dip for the *Nitrosopumilus* network and a peak for the *Sulfurovum* network between 30 and 50 mg/kg (Fig. 3I).

**Figure 3.**
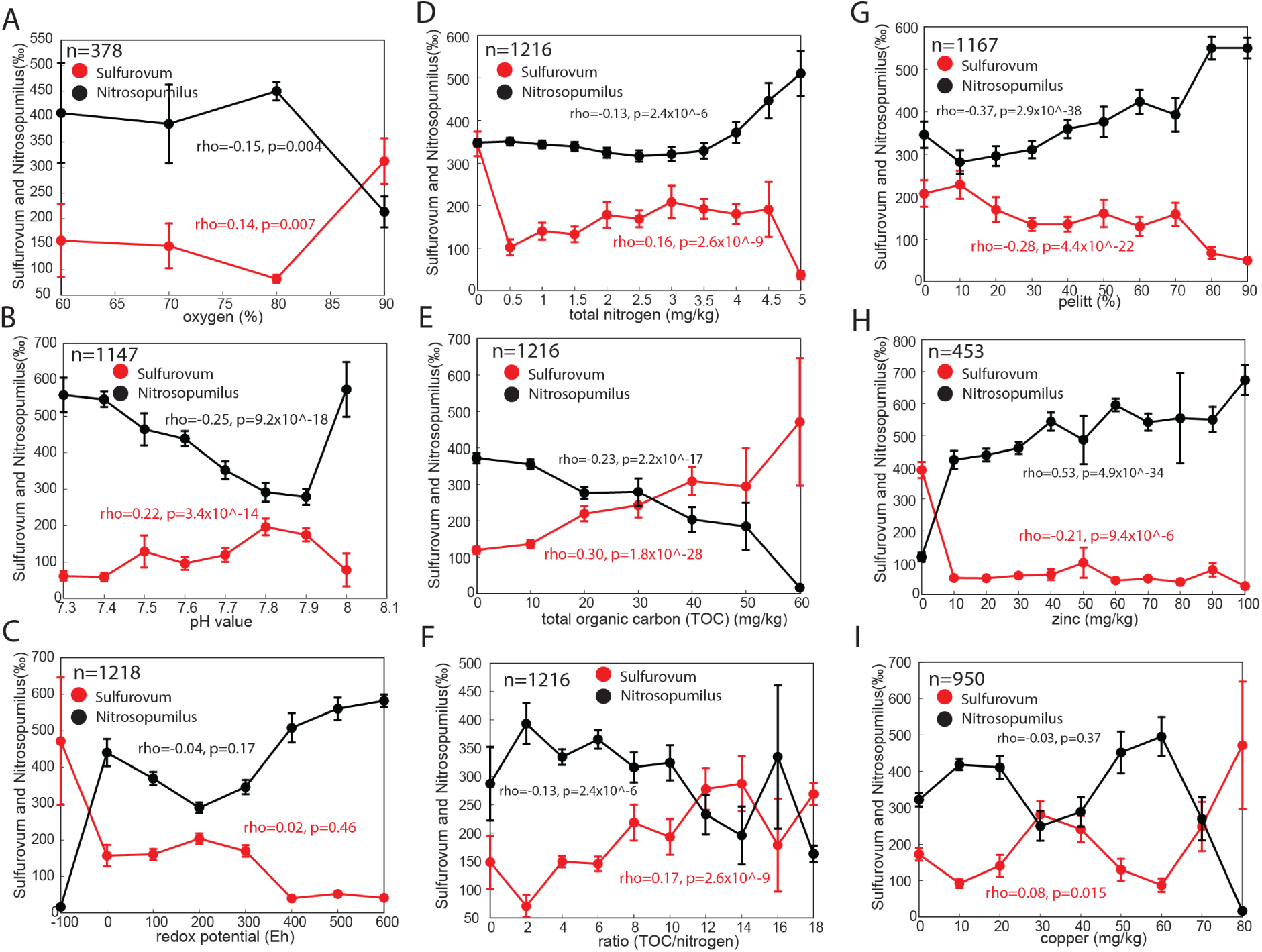
Chemical associations of the *Sulfurovum* and *Nitrosopumilus* networks. The panels illustrate the direct association between the *Sulfurovum* and *Nitrosopumilus* network clusters.

### Functional annotations showed distinct functionalities between the *Nitrosopumilus* and ***Sulfurovum* networks**

Functional investigations were conducted based on a catalogue of 357 metagenome assembled genomes (MAGs) obtained by shotgun sequencing of the same samples used for metabarcoding analysis above. Functional and taxonomic annotations are described in the Supplementary Information.

We selected 72 MAGs belonging to families unique to either the *Nitrosopumilus* (n=57 MAGs) or the *Sulfurovum* (n=15 MAGs) networks. Family level was used due to limited taxonomic assignments for marine species. Based on DRAM (Distilled and Refined Annotation of Metabolism) analyses we identified part of the 3-hydroxypropionate/4-hydroxybutyrate (3-HP/4-HB) cycle as overrepresented for the *Nitrosopumilus* network (Fig. 4A). For the *Sulfurovum* network, we identified MAGs classified as containing dissimilatory nitrate reduction to ammonium (DNRA) potential as overrepresented (Fig. 4B). In addition, we identified a significant effect for Complex IV low affinity Cytochrome C oxidase, with a mean value of 0.033 for the *Sulfurovum* network, and 0.12 for the *Nitrosopumilus* network (p=2.7×10^-4, FDR corrected Kruskal Wallis test). There was also a significant difference related to the potential for conversion of acetate to carbon dioxide and hydrogen. This was overrepresented in the *Sulfurovum* network with a prevalence of 60 % (9 out of 15), while the prevalence for the *Nitrosopumilus* network was 4 % (8 out of 57), p=3.8×10^-4, FDR corrected Chi-Square test.

**Figure 4.**
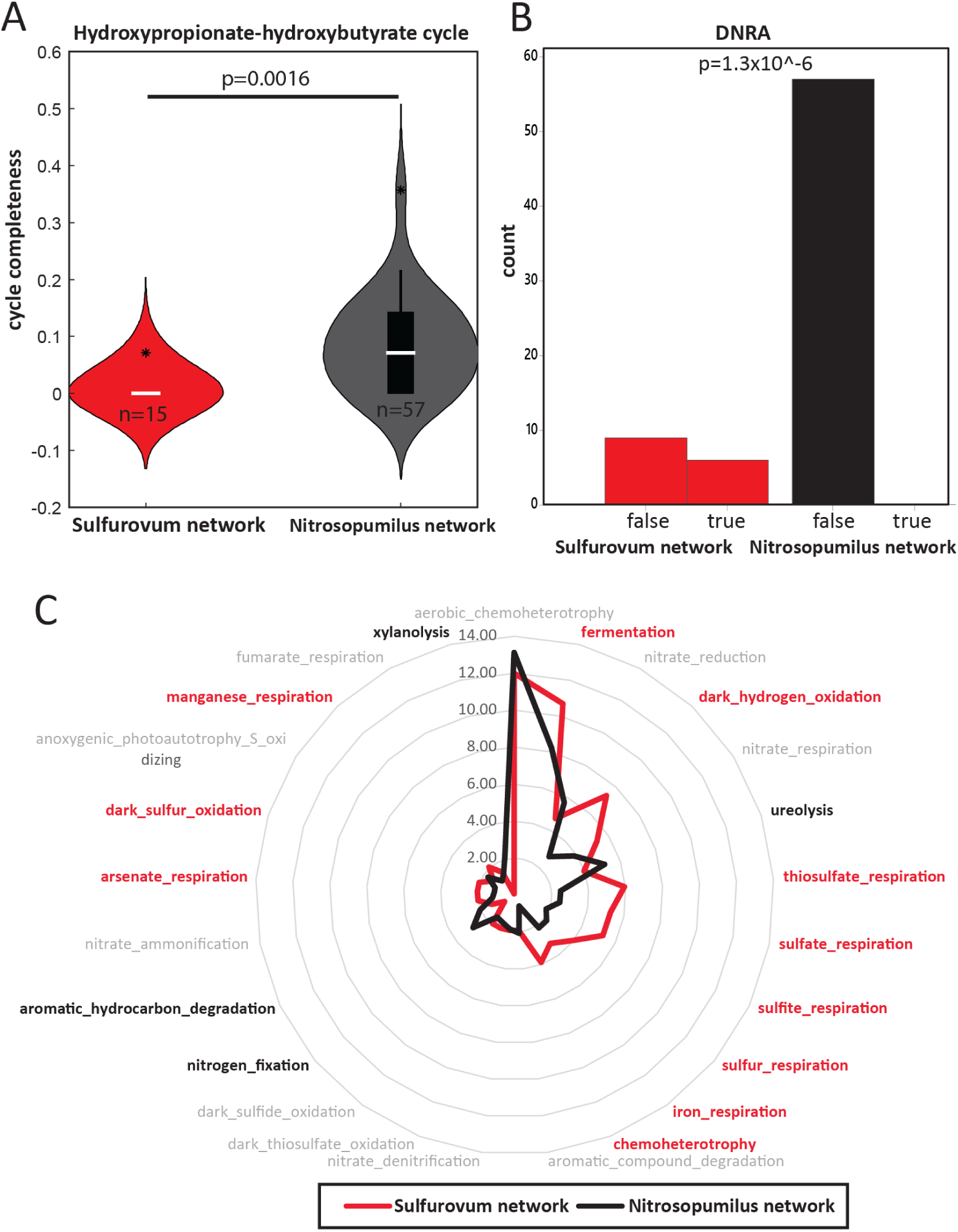
Functional properties associated with the *Sulfurovum* and the *Nitrosopumilus* network. (A) Fraction of metabolic cycle that was significantly different between the *Sulfurovum* and the *Nitrosopumilus* network, as determined by an FDR corrected Kruskal Wallis test of group abundances for functional annotations from DRAM. (B) Presence (true/false) of metabolic pathway, significantly differing between the *Sulfurovum -*, and the *Nitrosopumilus* network, as determined by FDR corrected Chi square analyses of categorical functional annotations from DRAM. (C) Illustration of the percentage of genera within the *Sulfurovum* and *Nitrosopumilus* network with the Faprotax functional assignment, as determined from 16S rRNA gene analyses. Functions covering more than 2 percent of the genera are included. Functions are assigned to the respective networks, given an absolute log10 ratio above 0.1.

As determined by the 16S rRNA gene-based Faprotax database^24^, the *Nitrosopumilus* network also exhibited a higher functional diversity than the *Sulfurovum* network, with 80 functions identified for the former, as compared to 36 for the latter (Suppl Table 1). For the most abundant functions (overall mean above 4 %), fermentation, hydrogen oxidation and sulfur respiration were overrepresented in the *Sulfurovum* network, while ureolysis, nitrogen fixation and aromatic hydrocarbon degradation were overrepresented for the *Nitrosopumilus* network (Fig. 4C). There was also a range of low-abundant traits that were overrepresented for the *Nitrosopumilus* network. These included one carbon metabolism, ammonium oxidation, cellulose and plastic degradation (Suppl Table 2).

### Bimodal distribution of *Nitrosopumilu*s and *Sulfurovum* networks

The *Nitrosopumilus* network displayed an apparent bimodal distribution pattern among sediment samples, with two overlapping peaks (Suppl. Fig. 2A), as evidenced by the Hartigan dip test of multimodality with a dip statistic of 0.052 and a p-value of 5.7×10^-4. A minimum value for the dip was reached at approximately ∼330 ‰. On the other hand, *Sulfurovum* exhibited a dip statistic of 0.06 and an p-value of less than 1×10^-5, with the first peak ending around ∼100 ‰ (Suppl. Fig 2B).

Based on the dip statistics, we converted the data to binary format at 330 ‰ for the *Nitrosopumilus* network, and at 100 ‰ for the *Sulfurovum* network. The binary data was then visualized in a PCoA plot based on Bray Curtis distances. This analysis revealed that the *Nitrosopumilus* network cluster is notably more compact compared to the *Sulfurovum* network cluster, which exhibited higher dispersion (Fig. 5A). The samples containing both the *Nitrosopumilus* and the *Sulfurovum* networks were located at the intersection between the two clusters along the first PCoA axis.

**Figure 5.**
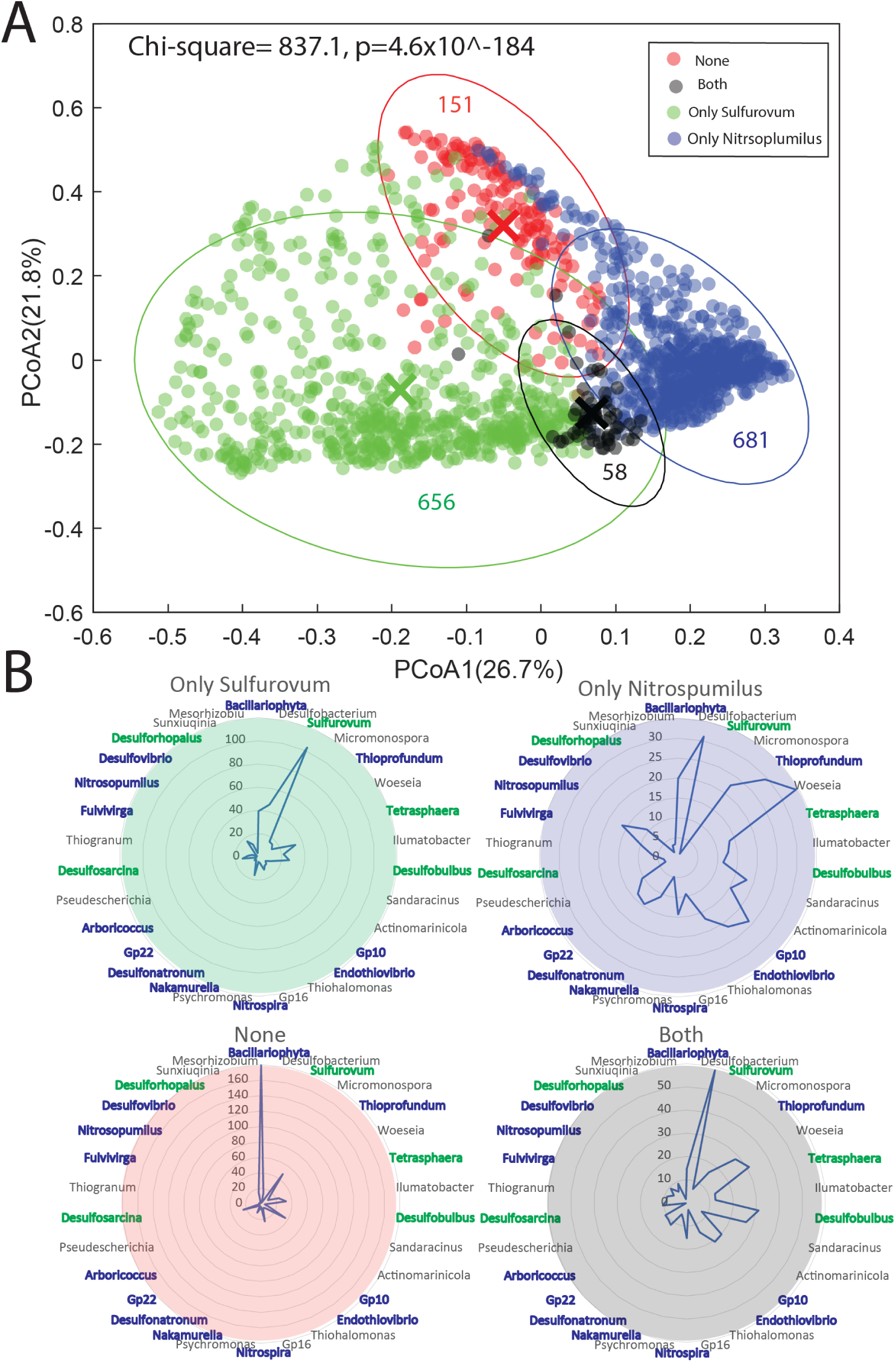
Taxonomic composition for the *Sulfurovum* and *Nitrosopumilus* network clusters. (A) PCoA plot illustrating for the overall beta diversity for the samples containing the *Sulfurovum* and *Nitrosopumilus* network clusters. (B) Mean distribution (‰) of the dominating genera within the different networks. The genera are color-coded with blue if they belong to the *Nitrosopumilus* network cluster, and green if they belong to the *Sulfurovum* network cluster. The ellipses represent 95% confidence interval, while the crosses represent the centroids.

The genus-level composition showed that *Sulfurovum* dominated in sediments whose prokaryote communities were dominated by OTUs associated with the *Sulfurovum* network. There was a larger range of genera that were observed to be abundant in sediments dominated by OTUs associated with the *Nitrosopumilus* network. Samples with ambiguous presence of both *Nitrosopumilus* and *Sulfurovum* networks showed a high relative abundance of chloroplasts from Bacillariophyta, while the intersect network contained a mix of *Sulfurovum*- and *Nitrosopumilus*-associated genera (Fig. 5B).

### Association of binarized data with ecological state and geography

To determine the risk of a mild deteriorated ecological state (nEQR < 0.8) for samples classified as rich in OTUs associated with either the *Nitrosopumilus* or the *Sulfurovum* networks, we calculated the respective odds ratios (OR). Sediments classified as rich in the *Sulfurovum* network had greater than a tenfold increase in risk of a deteriorated ecological state, with nEQR values below 0.8 (95 % CI: OR 9.5 to 15.6). In contrast, sediments classified as rich in the *Nitrosopumilus* network showed a tenfold decreased risk of ecological state deterioration (95 % CI: OR 0.068 to 0.11).

The geographical distribution showed a very clear east-west gradient. All the Islandic sediments were classified as rich in the *Sulfurovum* network, with no sediment samples having a particularly good ecological state with nEQR values above 0.8. On the other hand, for the samples from the North Sea, we found a higher prevalence of sediment samples classified as rich in the *Nitrosopumilus* network tcoincided with particularly good (nEQR > 0.8) ecological state classification. There was also a shift towards more sediment samples classified as *Sulfurovum*-enriched farther north in Norway (Fig. 6). The north to south gradient, however, is confounded by regional differences in investigation type.

**Figure 6.**
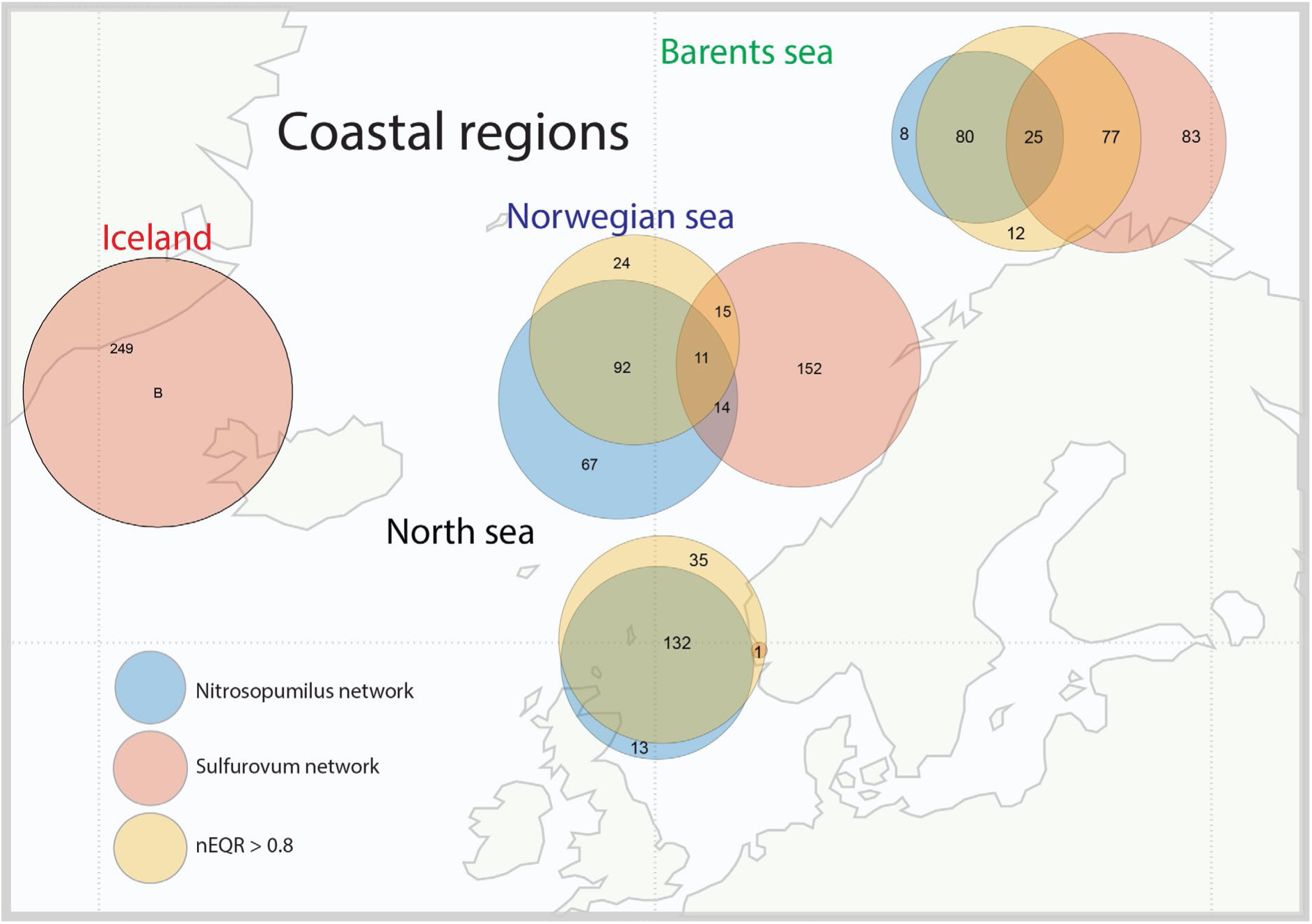
Geographical distribution of *Sulfurovum* and *Nitrosopumilus* network clusters. The Venn diagrams illustrate the association between the *Nitrosopumilus* and *Sulfurovum* network clusters and high nEQR values (> 0.8).

## DISCUSSION

The idea of keystone taxa, although controversial, underscores the significant ecological roles certain taxa play ^25,26^. In the context of the current study, *Nitrosopumilus* and *Sulfurovum* are candidates for potential keystone genera in the northern coastal sea regions investigated here. This is both due to their centrality in correlation networks ^27^ and the strong association with the macrofauna. In particular, *Nitrosopumilus* seems important as a keystone taxon in coordinating a large network of more than 500 positively correlated genera, being strongly associated with slightly to unperturbed ecological states based on macrofaunal assessments.

In our work, the association between the *Nitrosopumilus* network and the macrofauna demonstrated an apparent threshold, with relative abundance in metabarcoding data above ∼ 20% supporting a particularly good ecological state according to the GAM model. This may indicate that *Nitrosopumilus* provides essential components to the macrofauna. In the current work we identified both cobalamin dependent calcium carbonate fixation and cytochrome oxidase as associated with the *Nitrosopumilus* network. Previous research has suggested *Nitrosopumilus* as an essential contributor of cobalamin ^21,28,29^, with cobalamin being essential to maintain macroscopic life and support carbon dioxide fixation ^22^.

Cytochrome oxidase is a key enzyme in aerobic metabolism, indicating oxic conditions in the sediments associated with the *Nitrosopumilus* network ^30^. Additionally, *Nitrosopumilus* show strong correlations with over 500 other genera, likely offering other, yet unidentified, ecological services vital for the macrofauna in the sea.

From the GAM model, we found that the *Sulfurovum* network was associated with a gradual decline in macrofauna. From the MAGs we identified that dissimilatory nitrate reduction to ammonia (DNRA) was overrepresented for the *Sulfurovum* network. DNRA conserves ammonium on the seafloor ^31^, potentially leading to macrofauna deterioration because of ammonia toxicity ^32^. The combination of sulfides and DNRA has also been linked to nitrous oxide production ^33^. Although, *Sulfurovum* was positively associated with cytochrome oxidase, the rest of the genera in the *Sulfurovum* network showed negative association to cytochrome oxidase. This may indicate that *Sulfurovum* preserves a boundary between the oxic sea and anoxic sediments through sulfide oxidation. Furthermore, as for the *Nitrosopumilus* network, there are likely unknown mechanisms also associated with macrofauna ecological status for the *Sulfurovum* network.

For the energy yielding processes, our results support that the key processes are related to ammonium and sulfur oxidation, in addition to dismutation of acetate to carbon dioxide and hydrogen. The ammonium oxidation, however, seems mutually exclusive, with low overlap in samples containing genera that can perform sulfide oxidation and/or acetate dismutation. This result is in line with a recent finding by our group ^8^ as well as a recent finding that the redox chemistry of sediments and the oceans are bistable ^34^. Mechanistically, the competition between the *Sulfurovum* – and *Nitrosopumilus* networks could be related to sulfides disrupting cross-feeding between ammonium- and nitrite oxidizers, in addition to sulfide oxidation promoting nitrous oxide production ^35^. The sulfide driven oxidation by *Sulfurovum* could therefore outcompete the ammonium and nitrite oxidation processes driven by *Nitrosopumilus* ^10^. The potential increased nitrous oxide production could also interfere with the carbon dioxide fixation by archaeal ammonium oxidizers, due to the requirement for cobalamin in this process ^10^. Nitrous oxide inhibits cobalamin dependent metabolism ^36^. The potential release of hydrogen from acetate dismutation could contribute to sulfide generation through sulfate reduction, in addition contributing to high carbon dioxide levels, supporting autotrophic sulfide oxidizers ^37^.

We identified a strong east-west pattern with high *Sulfurovum* and low nEQR in the Icelandic samples. This pattern could be due to geothermal activity along the Atlantic Ridge, rather than human impact. Basalt from the Mid Atlantic Ridge contains elevated levels of sulfides ^38^, which could be a driver for sulfide oxidation. Suflide oxidation could also be the cause for the reduced ecological status observed around Iceland. However, there could also be other confounding factors, such as increased carbon dioxide and decreased pH ^39^.

The apparent overrepresentation of high nEQR for the North Sea coastal regions could potentially be due to a bias in sampling design, since samples were not collected from a representative balance of aquaculture impact. This bias could also be the reason for the overrepresentation of *Nitrosopumilus* in the North Sea region. Balanced sampling design, among other aspects, should be addressed in more targeted studies.

There was a surprising positive association between *Nitrosopumilus* and total nitrogen, while there was a negative association of this network with organic carbon. This may indicate a competitive advantage for *Nitrosopumilus* at low TOC:nitrogen ratios. A low ratio may prevent the outgrowth of heterotrophs since organic carbon is limiting. A high TOC:N ratio, on the other hand, promotes the *Sulfurovum* network. This is paradoxical, as *Sulfurovum* is also an autotroph. One potential explanation, however, could be that high TOC:N ratios promote the production of sulfides ^40^, which *Sulfurovum* could oxidize. This would sustain high oxygen usage and create a relentless cycle of sulfur reduction and oxidation ^11^.

We observed an opposite bimodal association of copper with the *Nitrosopumilus* and *Sulfurovum* networks. In seawater, copper is involved in complexing with sulfides ^41^, in addition to being essential for the reduction of nitrous oxide to nitrogen gas ^42^. The multifaceted interaction pattern of copper could be the reason for the bimodal association pattern towards *Sulfurovum* and *Nitrosopumilus*. Surprisingly, zinc showed a strong positive association with the *Nitrosopumilus* network. This association could be due to the dependence of carbonic anhydrase on zinc for catalysis of carbon dioxide to bicarbonate^43^. *Sulfurovum,* on the other hand, can utilize carbon dioxide directly without enzymatic conversion to bicarbonate ^37^. This may indicate that inhibition of bicarbonate formation through zinc limitation could provide a competitive advantage for *Sulfurovum*. Finally, we discovered a strong positive correlation between pelite and the *Nitrosopumilus* network.

This may indicate that *Nitrosopumilus* thrives in regions with weak bottom currents, thus representing potential hotspots for supporting macrofauna diversity ^44^. Seafloor sites with weak bottom currents may also be more vulnerable towards accumulation of waste deposits due to higher deposition rates in comparison with sites with strong bottom current ^45^.

Taken together, the relationships identified here potentially provides a meaningful, function-based relationship between microbial networks and macrofauna that can be used for the purpose of environmental monitoring and management in the future.

## MATERIALS AND METHODS

### Sample collection

The sediment samples were collected at 41 different sites along the Norwegian costal line which spans three main bodies of water, being the Barents Sea, Norwegian Sea and the North Sea, and 9 different sites along the western Icelandic coast were collected from 2021 to 2023. For the purposes of this study, samples which were collected in the coastal zones and within fjords, are referred to in three latitudinal groups according to those water bodies. Most of the samples were collected from aquaculture sites, but other samples were also included, such as those sampled prior to the establishment of aquaculture farms. An overview of the samples included is provided in Suppl Table 1.

From each station two Van Veen grabs were collected, one intended for chemical- and eDNA analyses, and the other for macrofauna. The macrofauna and chemical analyses were conducted according to requirements by Norwegian authorities ^19^, with the inclusion of sampling for eDNA from the grab intended for chemical analyses.

### eDNA sampling and extraction

Sample material was collected from the upper 1 cm of the undisturbed sediment surface by inserting a spoon through the inspection hatch of the grab. The sediment samples were frozen at −20 °C for shipment and further processing in the laboratory.

All eDNA extractions were done using a KingFisher Flex automated extraction platform (Thermo scientific, USA), with the magnetic bead based MagAttract PowerSoil KF kit (QIAGEN, Germany).

250 µL from each of the homogenized sediment samples were added to the bead plate, and we did a mechanical lysis for 4 x 30 sec at 1,800 rpm on a FastPrep 96 (MP Biomedicals, USA). All the centrifuge steps were done for 21 min at 1300 x g on a plate centrifuge, PlateSpin II (KUBOTA, Japan). The rest of the extraction was done following the manufacturers protocol.

Each plate contained a mock community, working as a positive control under the extraction and further processing of the samples.

### PCR amplification and 16S rRNA gene library preparation

The V3-V4 region of the 16S rRNA gene was amplified using 0.2 µM of the primers PRK341 forward (5’-CCTACGGGRBGCASCAG-3’) and PRK806 reverse (5’-GGACTACYVGGGTATCTAAT-3’) ^46^, together with 1x HOT FIREPol Blend Master Mix Ready to Load (Solis BioDyne, Estonia). The following amplification program was used: 95°C for 15 min, 25 cycles of 95°C for 30 sec, 55°C for 30 sec and 72°C for 45 sec, followed by 72°C for 7 min.

To purify the PCR products, we used 1x volume of Sera Mag Beads (Supplier). Amplicon purification was performed on Biomek 3000 or4000 (Beckman Coulter, USA) automated liquid handling platforms, following the manufacturers protocol. The program for PCR purification was the same on both instruments. Some plates were also done manually, due to technical issues, following the same protocol as on the instruments.

To index the amplicons, 1x FIREPol Master Mix Ready to Load (Solis BioDyne, Estonia) was used together with a combination of 16 forward and 36 reverse primers, with Illumina indexes. The following PCR program was used: 95°C for 5 min, 10 cycles of 95°C for 30 sec, 55°C for 1 min and 72°C for 45 sec, followed by 72°C for 7 min.

To concentration-normalize and pool all the samples to one library, we quantified the samples using a VarioSkan LUX Multimode Microplate Reader (ThermoFisher Scientiffic, USA) to measure the relative fluorescence unit (RFU) in each sample at 260 nm wavelength. We then made a standard curve from measured ng/µL using a Qubit fluorometer and the dsDNA High Sensitivity assay kit. This standard curve was used to estimate the concentration in ng/µL in each sample, which was used in further calculations for normalization. In order to get a deep sequencing, due to the high diversity of the sediment samples, each library consisted of approximately 288 samples. A purified library was sent to the Norwegian Sequencing Centre in Oslo, and sequenced on an Illumina MiSeq v3 platform.

### Whole genome shotgun sequencing

To perform a whole genome shotgun sequencing on a selection of 96 sediment samples, we used the Illumina Nextera DNA Flex Library Prep. The input of genomic DNA was between 10 and 49 ng, and we followed the manufacturers protocol according to the amount of DNA. The index adapters we used was the Nextera DNA UD Indexes, set C (IDT for illumina, USA), and during the library clean-up we followed the protocol for small PCR fragments (< 500 bp). The finished library was sequenced on a NovaSeq 6000 PE150 at Novogene UK.

### Data processing, assignments of taxonomy and function

The 16S raw data was demultiplexed and then processed using the VSEARCH software version 2.22.1^47^. Reads were trimmed (3’ end, 20 bases off R1 and 60 off R2 reads), merged and filtered by the fastq quality scores (mean error probability < 0.01). De-replicated reads were clustered into OTUs at 95% identity and checked for chimeras. The 95% level was chosen because of the large size dataset, and the high diversity of the samples. A table of read counts for all OTUs in all samples were computed. The OTU sequences were given a taxonomic classification using the SINTAX algorithm ^48^ implemented in VSEARCH, and with the RDP database ^49^. No threshold was used for the taxonomic assignment. Functional assignments were done using the FAPROTAV 1.2.7 database ^24^.

The reads from the whole genome sequencing were trimmed, filtered and merged (if possible) using the Bbmap software version 39.01 (sourceforge.net/projects/bbmap/). Each sample was assembled using SPAdes version 3.15.5 ^50^ in “meta” mode. Coverages for the resulting contigs in each sample were computed by coverM version 0.6.1 (github.com/wwood/CoverM) and the contigs were binned using both MaxBin2 version 2.2.7 ^51^ and MetaBat2 version 2.15 ^52^. All bins were given quality scores (completeness and contamination) with checkM2 version 1.0.1 ^53^ and then de-replicated with dRep version 3.4.0 ^54^. De-replicated bins with completeness at least 75% and contamination below 25% were considered the Metagenome Assembled Genomes (MAGs) and these were given a taxonomic assignment with GTDBTk version 2.3.2 ^55^ using the GTDB version 2.07 database. We also computed the mean coverage for all MAGs in all samples, again using coverM, and finally all MAGs were given functional annotations with DRAM version 1.4 ^56^.

### Statistical and ecological analyses

All statistical analyses were done using the implementations in the Matlab R2023 programming environment. We mostly used preprogrammed functions that can be run in Matlab. For the analyses involving pairwise correlations, we used the ‘corr’ function with Spearman correlation. Non-parametric tests were used to avoid linearity assumption. To determine differences between groups of continuous variables, we used the ‘kruskalwallis’ function, which implements the non-parametric Kruskal-Wallis test. The distribution of categorical variables was investigated using the Chi-Square Test of Independence; this test is implemented in the Matlab ‘crosstab’ function. All p-values were corrected for false discovery rate (FDR) using the ‘mafdr’ function with the Benjamini and Hochberg correction^57^.

Modelling of non-linear associations was done using cross validated Generalized Additive Models (GAM) with the function ‘fitgram’, using the parameters: ‘CrossVal’, ‘on’, ‘KFold’, 10, ‘FitStandardDeviation’, true. The function was also run without cross-validation for plotting purposes. The partial dependences were plotted using the ‘plotPartialDependence’ function, with the parameters: ‘Conditional’ and ‘absolute’. Due to the relatively large differences in the microbiota composition across all individual sediment samples ^58^, we did not take into account the hierarchical structure of the data in the GAM modelling.

The alpha diversity analyses were done using an in-house Matlab implementation of the Simpson’s D and Shannon H indexes. For the 1 - Simpson’s D index we used the following implementation: 1-D = 1 **-** ∑(*pi*)^2^, where *pi* is the portion of a specific OTU. For Shannon H we used the following implementation: H=−∑(*pi* ×ln*pi*). The beta-diversity analyses were conducted using the Matlab Fathom package (www.usf.edu/marine-science/research/matlab-resources/fathom-toolbox-for-matlab.aspx). A Bray Curtis distance matrix was generated using the Fathom function ‘f_dis’ with the ‘bc’ parameter. The ‘f_dis’ distance matrix was the used as input in the ‘f_pcoa’ function, which performs a Principal Coordinates Analysis (PCoA). The output from the ‘f_pcoa’ function was finally used as an input in the ‘f_pcoaPlot’ function, generating a graphical representation of the PCoA plot. Binarization of the 16S rRNA gene sequence data was based ‘hartigansdipsigniftest’, with 10 000 bootstrap replicates. This test determines the deviation from a unimodal distribution, as outlined by Hartigan ^59^. If a statistically significant deviation from unimodality was identified (p<0.05), then a threshold was set to binarize the data.

The odds ratios were determined, assuming that nEQR values below 0.8 represent deteriorated ecological states, while values above 0.8 represented healthy states. The exposure was represented by the presence of the respective groups of microorganisms. The ratios were calculated dividing the relative occurrence of macrofauna loss in the presence of the respective group of microorganisms on the relative occurrence in the absence of the group of microorganisms. The 95% confidence intervals were calculated as described by Altman ^60^.

## Data availability

The raw 16S rRNA gene sequencing data are available under the accession number PRJNA1128851 in the Sequence Read Archive (SRA).

## Code Availability

All code for the raw data processing is found in the compressed archive at https://arken.nmbu.no/~larssn/nitrosulf.tar.gz. This archive contains a subfolder for code used for processing the 16S data, another subfolder for code used for processing WGS data, and a third subfolder with apptainer containers. Scripting was done in R or UNIX shell and any external software tools used are supplied as apptainer containers. All processing of rawdata was done on a High Performance Computing cluster using a Centos Linux 7.9 operating system.

## SUPPLEMENTARY INFORMATION

**Suppl Figure 1.**
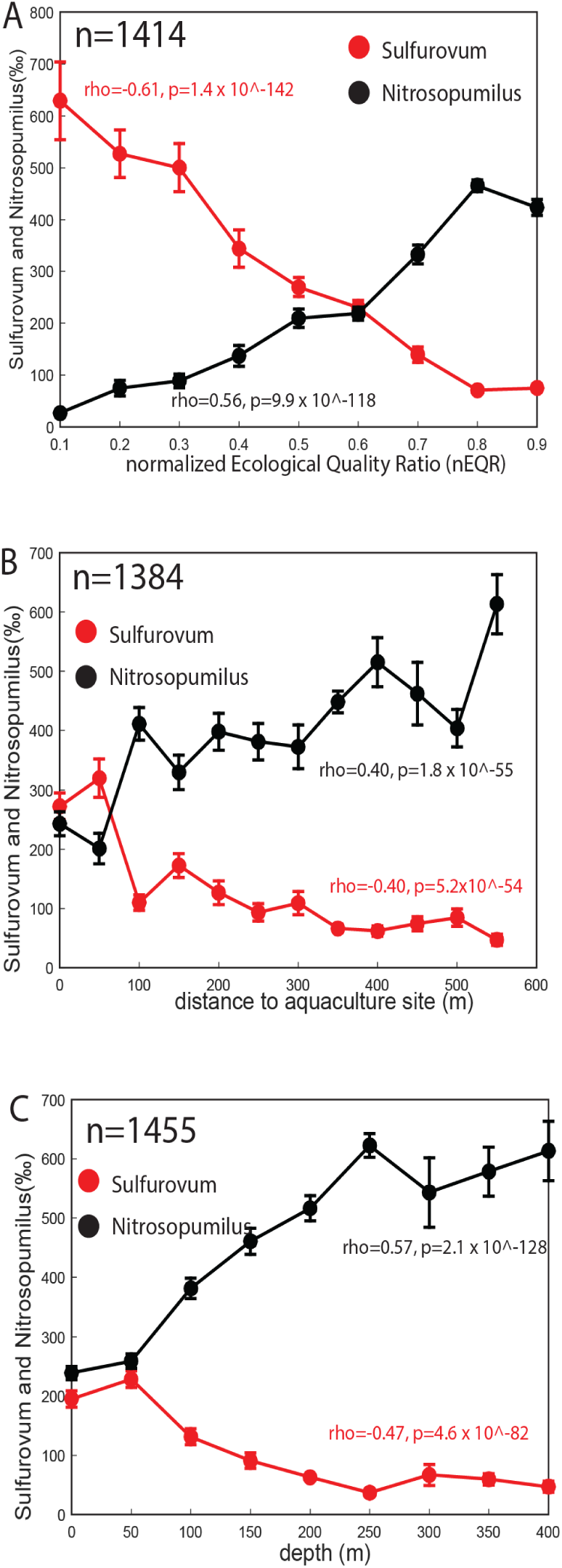
Association between *Sulfurovum* and *Nitrosopumilus* networks. The panels illustrate the direct association between the Sulfurovum - and Nitrosopumilus networks.

**Suppl Figure 2.**
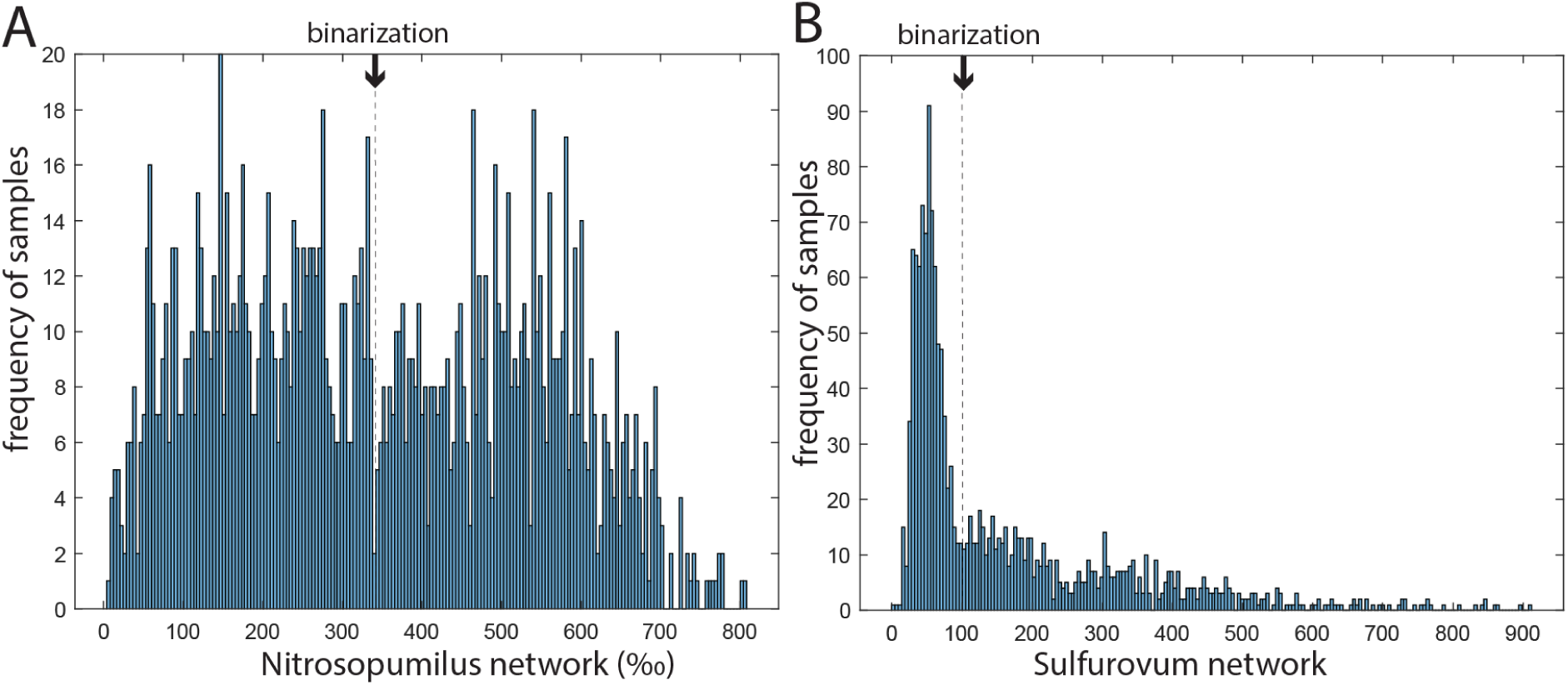
Frequency of (A) the *Nitrosopumilus* - and (B) the *Sulfurovum* network. The thresholds for binarization to the respective network clusters are marked.

---

**Suppl Table 1.** Sampling locations.

| Country | Water body<sup>1</sup> | Examination | # Locations<sup>2</sup> | # Samples<sup>3</sup> |
| --- | --- | --- | --- | --- |
| Norway | Norwegian Sea | Pre examination | 5 | 165 |
| Norway | Norwegian Sea | ASC-examination | 3 | 96 |
| Norway | Norwegian Sea | C-examination | 5 | 210 |
| Norway | Norwegian Sea | Recipient examination | 1 | 60 |
| Norway | Barents Sea | Pre examination | 1 | 36 |
| Norway | Barents Sea | C-examination | 7 | 237 |
| Norway | North Sea | Pre examination | 11 | 312 |
| Norway | North Sea | C-examination | 2 | 30 |
| Norway | North Sea | Recipient examination | 2 | 12 |
| Iceland | Danish Strait | Pre examination | 1 | 30 |
| Iceland | Danish Strait | C-examination | 8 | 244 |

<sup>1</sup>Water body is defined as those with longitude below 0°east – Danish Strait, latitude below 60° north – North Sea, between 60° and 70° north – Norwegian Sea, and above 70° north – Barents sea.
<sup>2</sup>Four locations lacked geocoordinates. <sup>3</sup>One hundred and fourteen samples lacked geocoordinates.

**Suppl Table 2.**
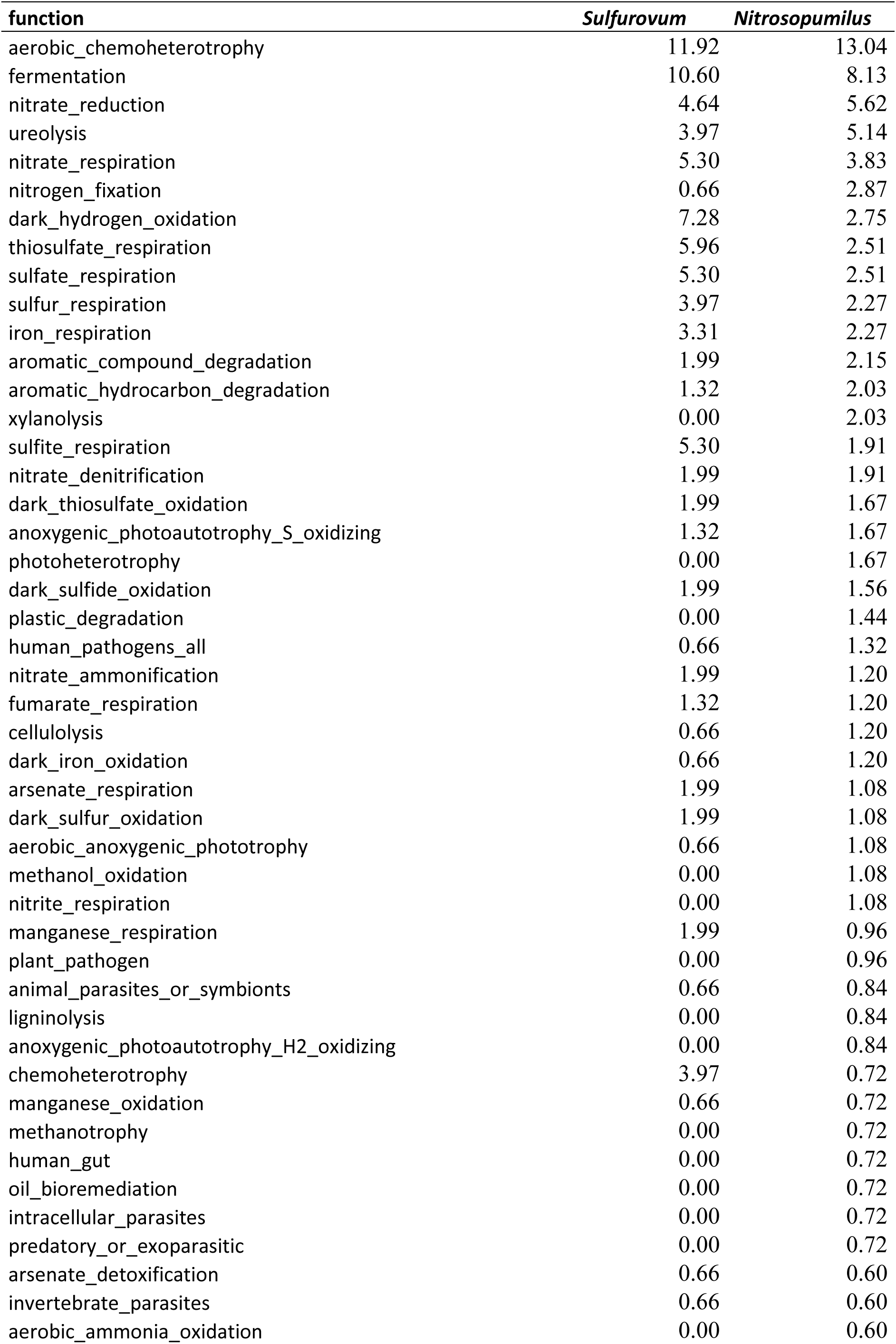
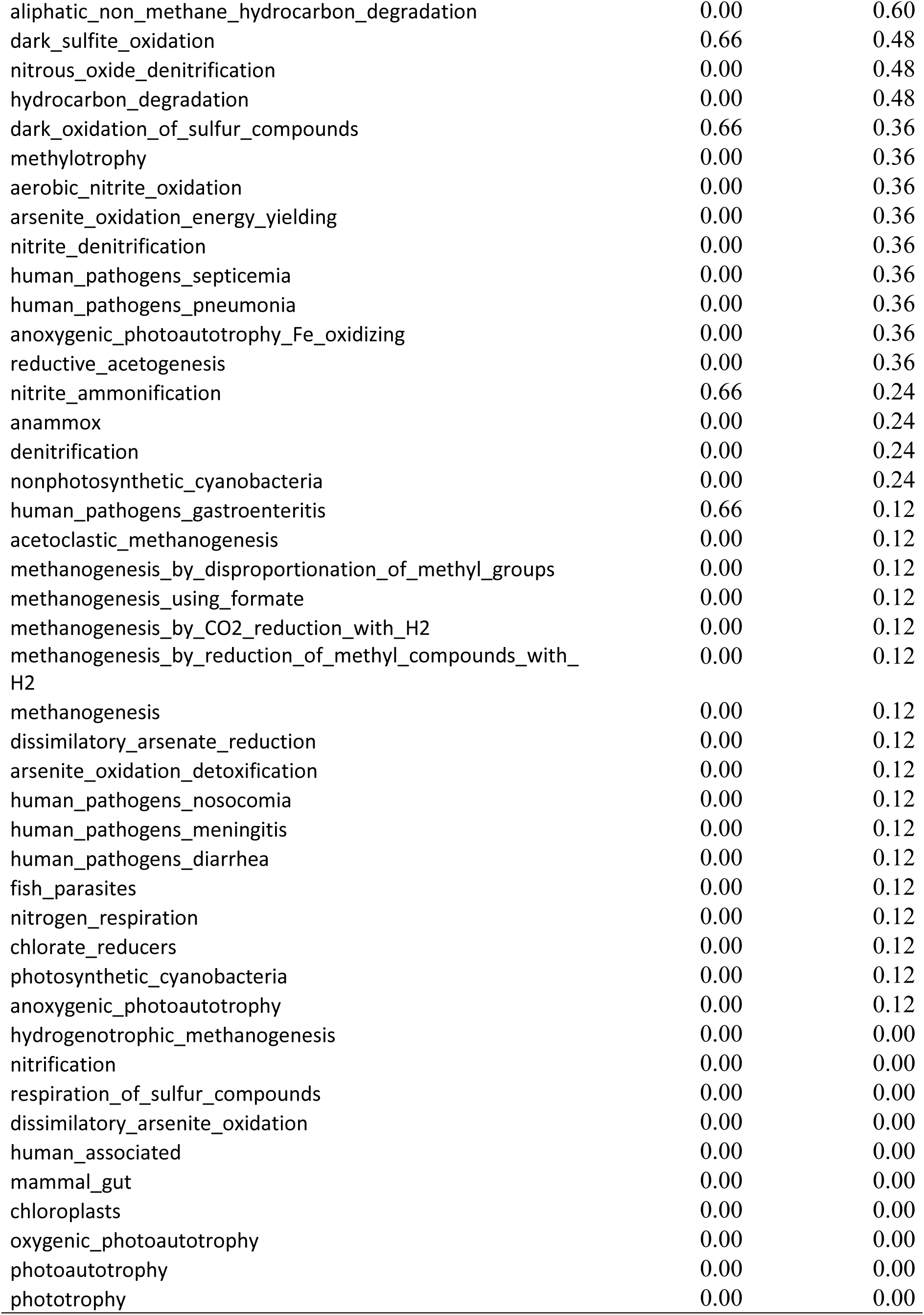
Functional assignment with the Faprotax database (percentage)

## REFERENCES

1. Snelgrove, P.V.R. Getting to the Bottom of Marine Biodiversity: Sedimentary Habitats: Ocean bottoms are the most widespread habitat on Earth and support high biodiversity and key ecosystem services. BioScience 49, 129–138 (1999).

2. Orsi, W.D. Ecology and evolution of seafloor and subseafloor microbial communities. Nature Reviews Microbiology 16, 671–683 (2018).

3. Laiolo, E. et al. Metagenomic probing toward an atlas of the taxonomic and metabolic foundations of the global ocean genome. Frontiers in Science 1(2024).

4. Clinton, M.E., Snelgrove, P.V.R. & Bates, A.E. Macrofaunal diversity patterns in coastal marine sediments: re-examining common metrics and methods. Marine Ecology Progress Series 735, 1–26 (2024).

5. Keeley, N. et al. Resilience of dynamic coastal benthic ecosystems in response to large-scale finfish farming. Aquaculture Environment Interactions 11(2019).

6. Besley, C.H. & Birch, G.F. Deepwater ocean outfalls: A sustainable solution for sewage discharge for mega-coastal cities (Sydney, Australia): Influence of deepwater ocean outfalls on shelf benthic infauna. Mar Pollut Bull 145(2019).

7. Ramirez-Llodra, E. et al. Submarine and deep-sea mine tailing placements: A review of current practices, environmental issues, natural analogs and knowledge gaps in Norway and internationally. Marine Pollution Bulletin 97, 13–35 (2015).

8. Pettersen, R. et al. Bimodal distribution of seafloor microbiota diversity and function are associated with marine aquaculture. Marine Genomics 66, 100991 (2022).

9. Könneke, M. et al. Isolation of an autotrophic ammonia-oxidizing marine archaeon. Nature 437, 543–546 (2005).

10. Walker, C.B. et al. <em>Nitrosopumilus maritimus</em> genome reveals unique mechanisms for nitrification and autotrophy in globally distributed marine crenarchaea. Proceedings of the National Academy of Sciences 107, 8818–8823 (2010).

11. Campbell, B.J., Engel, A.S., Porter, M.L. & Takai, K. The versatile epsilon-proteobacteria: key players in sulphidic habitats. Nat Rev Microbiol 4, 458–68 (2006).

12. Mahmood, M. et al. Increase in sedimentary organic carbon with a change from hypoxic to oxic conditions. Mar Pollut Bull 168, 112397 (2021).

13. Nascimento, J.R. et al. Microbial community shift under exposure of dredged sediments from a eutrophic bay. Environ Monit Assess 192, 539 (2020).

14. Olaussen, J.O. Environmental problems and regulation in the aquaculture industry. Insights from Norway. Marine Policy 98, 158–163 (2018).

15. Keeley, N., Wood, S.A. & Pochon, X. Development and preliminary validation of a multi-trophic metabarcoding biotic index for monitoring benthic organic enrichment. Ecological Indicators 85, 1044–1057 (2018).

16. Borja, A., Franco, J. & Pérez, V. A Marine Biotic Index to Establish the Ecological Quality of Soft-Bottom Benthos Within European Estuarine and Coastal Environments. Marine Pollution Bulletin 40, 1100–1114 (2000).

17. Poikane, S. et al. European aquatic ecological assessment methods: A critical review of their sensitivity to key pressures. Science of The Total Environment 740, 140075 (2020).

18. Andersen, J.H. et al. Approaches for integrated assessment of ecological and eutrophication status of surface waters in Nordic Countries. Ambio 45, 681–91 (2016).

19. Iversen, A. & Sandøy, S. Veileder 02:2018 Klassifisering av miljøtilstand i vann. Økologisk og kjemisk klassifiseringssystem for kystvann, grunnvann, innsjøer og elver. (ed. Environment, t.D.f.t.) (2018).

20. Hargrave, B.T., Holmer, M. & Newcombe, C.P. Towards a classification of organic enrichment in marine sediments based on biogeochemical indicators. Mar Pollut Bull 56, 810–24 (2008).

21. Doxey, A.C., Kurtz, D.A., Lynch, M.D., Sauder, L.A. & Neufeld, J.D. Aquatic metagenomes implicate Thaumarchaeota in global cobalamin production. Isme j 9, 461–71 (2015).

22. Sañudo-Wilhelmy, S.A. et al. Multiple B-vitamin depletion in large areas of the coastal ocean. Proceedings of the National Academy of Sciences 109, 14041–14045 (2012).

23. Balk, L. et al. Wild birds of declining European species are dying from a thiamine deficiency syndrome. Proceedings of the National Academy of Sciences 106, 12001–12006 (2009).

24. Louca, S., Parfrey, L.W. & Doebeli, M. Decoupling function and taxonomy in the global ocean microbiome. Science 353, 1272–1277 (2016).

25. Paine, R.T. A Note on Trophic Complexity and Community Stability. The American Naturalist 103, 91–93 (1969).

26. Mills, L., Soulé, M. & Doak, D. The Keystone-Species Concept in Ecology and Conservation. BioScience 43, 219–224 (1993).

27. Jiang, L. & Zhang, W. Determination of keystone species in CSM food web: A topological analysis of network structure. Network Biology 5, 13–33 (2015).

28. Beauvais, M. et al. Functional redundancy of seasonal vitamin B12 biosynthesis pathways in coastal marine microbial communities. Environmental Microbiology n/a.

29. Elling, F.J. et al. Vitamin B12-dependent biosynthesis ties amplified 2-methylhopanoid production during oceanic anoxic events to nitrification. Proc Natl Acad Sci U S A 117, 32996–33004 (2020).

30. Castresana, J., Lübben, M., Saraste, M. & Higgins, D.G. Evolution of cytochrome oxidase, an enzyme older than atmospheric oxygen. Embo j 13, 2516–25 (1994).

31. Hutchins, D.A. & Capone, D.G. The marine nitrogen cycle: new developments and global change. Nature Reviews Microbiology 20, 401–414 (2022).

32. Batley, G.E. & Simpson, S.L. Development of guidelines for ammonia in estuarine and marine water systems. Mar Pollut Bull 58, 1472–6 (2009).

33. Senga, Y., Mochida, K., Fukumori, R., Okamoto, N. & Seike, Y. N2O accumulation in estuarine and coastal sediments: The influence of H2S on dissimilatory nitrate reduction. Estuarine, Coastal and Shelf Science 67, 231–238 (2006).

34. van de Velde, S.J., Reinhard, C.T., Ridgwell, A. & Meysman, F.J.R. Bistability in the redox chemistry of sediments and oceans. Proceedings of the National Academy of Sciences 117, 33043–33050 (2020).

35. Delgado Vela, J., Bristow, L.A., Marchant, H.K., Love, N.G. & Dick, G.J. Sulfide alters microbial functional potential in a methane and nitrogen cycling biofilm reactor. Environmental Microbiology 23, 1481–1495 (2021).

36. Alston, T.A. Inhibition of vitamin B12-dependent microbial growth by nitrous oxide. Life Sci 48, 1591–5 (1991).

37. Kwon, H.-S. et al. Biofixation of a high-concentration of carbon dioxide using a deep-sea bacterium: Sulfurovum lithotrophicum 42BKTT. RSC Advances 5, 7151–7159 (2015).

38. Reekie, C.D.J. et al. Sulfide resorption during crustal ascent and degassing of oceanic plateau basalts. Nature Communications 10, 82 (2019).

39. Thomas, D.L., Bird, D.K., Arnórsson, S. & Maher, K. Geochemistry of CO2-rich waters in Iceland. Chemical Geology 444, 158–179 (2016).

40. Valdemarsen, T., Kristensen, E. & Holmer, M. Sulfur, carbon, and nitrogen cycling in faunated marine sediments impacted by repeated organic enrichment. Marine Ecology Progress Series 400, 37–53 (2010).

41. Al-Farawati, R. & van den Berg, C.M.G. Metal–sulfide complexation in seawater. Marine Chemistry 63, 331–352 (1999).

42. Sullivan, M.J., Gates, A.J., Appia-Ayme, C., Rowley, G. & Richardson, D.J. Copper control of bacterial nitrous oxide emission and its impact on vitamin B12-dependent metabolism. Proceedings of the National Academy of Sciences of the United States of America 110, 19926–19931 (2013).

43. Smith, K.S., Jakubzick, C., Whittam, T.S. & Ferry, J.G. Carbonic anhydrase is an ancient enzyme widespread in prokaryotes. Proceedings of the National Academy of Sciences 96, 15184–15189 (1999).

44. Guerra, C.A. et al. Global hotspots for soil nature conservation. Nature (2022).

45. Keeley, N.B., Forrest, B.M. & Macleod, C.K. Novel observations of benthic enrichment in contrasting flow regimes with implications for marine farm monitoring and management. Mar Pollut Bull 66, 105–16 (2013).

46. Yu, Y., Lee, C., Kim, J. & Hwang, S. Group-specific primer and probe sets to detect methanogenic communities using quantitative real-time polymerase chain reaction. Biotechnology and Bioengineering 89, 670–679 (2005).

47. Rognes, T., Flouri, T., Nichols, B., Quince, C. & Mahe, F. VSEARCH: a versatile open source tool for metagenomics. PeerJ 4, e2584 (2016).

48. Edgar, R.C. SINTAX: a simple non-Bayesian taxonomy classifier for 16S and ITS sequences. bioRxiv, 074161 (2016).

49. Wang, Q. & Cole, J.R. Updated RDP taxonomy and RDP Classifier for more accurate taxonomic classification. Microbiology Resource Announcements 0, e01063–23.

50. Nurk, S., Meleshko, D., Korobeynikov, A. & Pevzner, P.A. metaSPAdes: a new versatile metagenomic assembler. Genome Res 27, 824–834 (2017).

51. Wu, Y.-W., Simmons, B.A. & Singer, S.W. MaxBin 2.0: an automated binning algorithm to recover genomes from multiple metagenomic datasets. Bioinformatics 32, 605–607 (2015).

52. Kang, D.D., Froula, J., Egan, R. & Wang, Z. MetaBAT, an efficient tool for accurately reconstructing single genomes from complex microbial communities. PeerJ 3, e1165 (2015).

53. Chklovski, A., Parks, D.H., Woodcroft, B.J. & Tyson, G.W. CheckM2: a rapid, scalable and accurate tool for assessing microbial genome quality using machine learning. *bioRxiv*, 2022.07.11.499243 (2022).

54. Olm, M.R., Brown, C.T., Brooks, B. & Banfield, J.F. dRep: a tool for fast and accurate genomic comparisons that enables improved genome recovery from metagenomes through de-replication. The ISME Journal 11, 2864–2868 (2017).

55. Chaumeil, P.-A., Mussig, A.J., Hugenholtz, P. & Parks, D.H. GTDB-Tk: a toolkit to classify genomes with the Genome Taxonomy Database. Bioinformatics 36, 1925–1927 (2019).

56. Shaffer, M. et al. DRAM for distilling microbial metabolism to automate the curation of microbiome function. Nucleic Acids Res 48, 8883–8900 (2020).

57. Benjamini, Y. & Hochberg, J. Controlling the false discovery rate: a practical and powerful approach to multiple testing. J R Stat Soc Ser B 57(1995).

58. Nilsen, T. et al. Swarm and UNOISE outperform DADA2 and Deblur for denoising high-diversity marine seafloor samples. ISME Communications (2024).

59. Hartigan, J.A. & Hartigan, P.M. The Dip Test of Unimodality. The Annals of Statistics 13, 70–84 (1985).

60. Altman, D. Practical Statistics for Medical Research, (Chapman and Hall/CRC, 1991).

